# Dibenzocyclooctyne-modified PCR primers enable direct enzyme-free click chemistry ligation for custom nanopore amplicon sequencing

**DOI:** 10.64898/2026.04.18.719403

**Authors:** Patrick Łypaczewski, B. Jesse Shapiro

## Abstract

Oxford Nanopore Technologies (ONT) rapid library preparation kits use transposase-mediated tagmentation to attach click chemistry functionalized oligonucleotide duplexes to fragmented DNA, followed by click chemistry to conjugate Rapid Adapter (RA) sequencing adapters. A similar protocol is used in 16S rRNA gene amplicon and PCR-amplified rapid whole-genome sequencing workflows. Here, we describe custom oligonucleotides with dibenzocyclooctyne (DBCO) added onto PCR primer 5′ termini. After standard PCR amplification, DBCO-modified amplicons react spontaneously with RA sequencing adapters, producing sequencing-ready libraries in minutes without enzymatic processing. All configurations employ an asymmetric design in which the DBCO modification is restricted to a single primer, leaving the opposite primer available for barcoding at low cost. We validate three primer architectures: (i) direct attachment of DBCO to a target-specific primer, (ii) a universal DBCO-modified oligonucleotide used in a two-step PCR workflow, and (iii) a three-primer single-pot reaction combining the universal DBCO oligonucleotide with unmodified target-specific primers.

These configurations are validated using full-length 16S rRNA gene amplicons sequenced on a PromethION flow cell. DBCO-modified primers are synthesized either commercially or in-house via DBCO-TFP ester conjugation to 5′-amino oligonucleotides and remain fully active through standard PCR thermocycling. The best-performing configuration used a two-step PCR with a universal oligonucleotide and achieved higher pore occupation and reads than comparable commercial solutions. This approach reduces library preparation reagent costs compared to available kits, as the initial synthesis cost is lower than existing amplicon sequencing kits, while providing enough material for hundreds or thousands of PCR reactions. This is further applicable to an unlimited number of gene targets beyond 16S sequencing.

## INTRODUCTION

Amplicon sequencing on Oxford Nanopore Technologies (ONT) platforms has become a cornerstone of microbial community profiling, viral genomic surveillance, and clinical diagnostics (Benítez-Páez et al., 2016; Charalampous et al., 2019; Quick et al., 2017). The ability to sequence full-length 16S rRNA genes (~1,500 bp), complete viral genomes via tiling amplicons, and targeted pathogen panels with real-time data acquisition distinguishes nanopore from short-read platforms (Calus et al., 2018; Freed et al., 2020). However, current ONT amplicon workflows impose a fundamental trade-off between simplicity and flexibility that limits their adoption in diverse research and clinical settings.

ONT’s bundled amplicon kits, the 16S Barcoding Kit (SQK-16S114.24) and the newer Microbial Amplicon Barcoding Kit (SQK-MAB114.24), offer streamlined, single-box workflows but carry substantial constraints. The 16S Barcoding Kit uses fixed 27F/1492R primers that harbour three mismatches against *Bifidobacterium* spp., causing complete amplification failure for this clinically important genus (Kai et al., 2019; Matsuo et al., 2021; Nygaard et al., 2020; Winand et al., 2020). This primer bias extends to *Borrelia*, Chlamydiales, *Gardnerella*, and Archaea, as independently confirmed across gut (Winand et al., 2020), oropharyngeal (Kai et al., 2019), and mock community samples (Loit et al., 2019; Nygaard et al., 2020). The kit is restricted to 24 barcodes, locked to 16S targets exclusively, has a reported 3-month shelf life at −20°C, contains limited primer volumes prohibiting troubleshooting and optimization reactions, and is scheduled for discontinuation in mid-2026. Users cannot modify the primer sequences, substitute alternative targets, or incorporate modifications such as unique molecular identifiers (UMIs) to increase single-read accuracy.

The SQK-MAB114.24 kit, released in September 2025 as an Early Access product, partially addresses primer bias by offering redesigned 16S primers with improved inclusivity, while adding ITS primers for fungal profiling and decoupling primers from the barcoding step (Oxford Nanopore, 2025). However, this kit introduces a multi-step post-PCR protocol that substantially offsets its flexibility gains: amplicons must undergo an enzymatic barcode ligation, followed by proteinase K digestion of barcoding enzymes, heat inactivation of the proteinase K, bead cleanup, and finally RA adapter attachment. This series of incubations and enzymatic steps extends hands-on time well beyond the simpler 16S Barcoding Kit workflow, and the kit retains the 24-barcode ceiling. The Rapid PCR Barcoding Kit (SQK-RPB114.24) offers an alternative four-primer approach with 24 barcodes but provides only a few microliters of 10 μM barcoded primers per barcode, a quantity that is rapidly exhausted in routine use, particularly for studies requiring optimization or replicate runs.

The general-purpose library preparation kits, ligation (SQK-LSK114) and rapid barcoding (SQK-RBK114.24 or SQK-RBK114.96), provide more flexibility but require extensive post-PCR processing. Ligation demands end-repair, dA-tailing, enzymatic adapter ligation, and two bead cleanups (~75 minutes). Rapid barcoding employs a MuA transposase that simultaneously fragments DNA and attaches functionalized oligonucleotide duplexes, followed by conjugation of the Rapid Adapter (Green et al., 2012; Haapa-Paananen et al., 2002; Picelli et al., 2014). While originally developed for whole-genome sequencing where this kit allows sequencing-ready libraries to be generated rapidly through one-pot fragmentation and barcoding combined with click chemistry, its adaptation to amplicon sequencing solves the problem of limited amplicon range but is not well-suited to non-clonal amplicons: the transposase cleaves within amplicons to generate sequencing reads in opposite directions, unlinking upstream and downstream sequence information. This “middle-out” reading additionally often leads to truncation of 10-20 bp from the amplicon ends as the DNA slips off the adapter and into the pore. The transposase also exhibits modest insertion site bias at 5′-TATGA motifs (Haapa-Paananen et al., 2002), and the proprietary MuA-loaded transposome reagent dominates per-sample costs.

Click chemistry has been applied to sequencing library construction in other contexts, most notably ClickSeq, which uses copper-catalyzed azide-alkyne cycloaddition (CuAAC) to ligate Illumina adapters to 3′-azido-terminated cDNA fragments (Jaworski & Routh, 2017; Routh, 2020; Routh et al., 2015). Schonegger et al. demonstrated CuAAC-based cDNA circularization for full-length transcript sequencing on ONT, though adapter attachment remained enzymatic (Schönegger et al., 2022). We decided against copper-mediated reactions as the effect of additional copper ions on the sensitive flow cell chemistry is unknown. SPAAC has been used for terminal DNA labelling (Marks et al., 2011), triazole-linked sequencing library construction (Stead et al., 2018), and NAD+-capped transcript enrichment (Hu et al., 2021). SPAAC is a copper-free bioorthogonal reaction between two partners: a small, linear azide (or tetrazide) and a strained cyclooctyne ring system. Critically, Marks et al. demonstrated that cyclooctyne-modified primers survive PCR thermocycling and remain reactive toward azide conjugation partners (Marks et al., 2011), establishing that modifications on primer 5′ termini are compatible with standard amplification protocols. Furthermore, copper-free conjugation moieties are readily available as modifications from commercial suppliers such as Integrated DNA Technologies (IDT).

We investigated the two modifications currently available for SPAAC at IDT: azide and DBCO. We reasoned that incorporating a modification group directly onto PCR primer 5′ termini would enable SPAAC conjugation with a complementary modification-bearing sequencing adapter immediately after PCR, eliminating all post-amplification enzymatic steps. Unlike the bundled kits, this approach imposes no constraints on primer sequence, target locus, barcode number, or primer modification. Unlike the MAB kit’s multi-step protocol, the entire post-PCR library preparation consists of a single cleanup and room-temperature RA adapter addition. And unlike the Rapid PCR Barcoding Kit, modified primers synthesized commercially typically yield 50-200 μL at 100 μM, sufficient for thousands of reactions from a single synthesis, compared to the limited 10 μM aliquots provided in commercial kits. Here, we describe the design, synthesis, optimization, and validation of DBCO-modified primers across multiple workflow configurations, benchmarked against standard ONT library preparation on PromethION flow cells using full-length 16S rRNA sequencing on a defined mock community.

## MATERIALS AND METHODS

### PCR amplification

All PCR reactions targeted the full-length 16S rRNA gene (~1,500 bp) using the ZymoBIOMICS Microbial Community DNA Standard (D6306, Zymo Research) diluted to 5 ng/μL as template. Amplification was performed using Platinum SuperFi II DNA Mastermix (Thermo Fisher Scientific) in a final reaction volume of 50 μL with 0.2 μM of each primer. Standard thermocycling conditions consisted of an initial denaturation at 98 °C for 30 seconds, followed by 45 cycles of 98 °C for 10 seconds, 60 °C for 10 seconds, and 72 °C for 45 seconds, with a final extension at 72 °C for 5 minutes. Second-round PCR reactions (where applicable) used 15 cycles with 1 μL of purified first-round product as template. Where different first round templates were used for second round PCR, the purified first round products were first diluted to the lowest concentration to maintain an even number of initial templates. All PCR products were verified by agarose gel electrophoresis prior to sequencing. Products were cleaned using a 0.5X ratio of AMPure XP beads (Beckman Coulter) and quantified on a Qubit fluorometer (Thermo Fisher Scientific).

### Oligonucleotide design and synthesis

The core primer pair consisted of 27F (5′-AGRGTTYGATYMTGGCTCAG-3’) and 1492R (5′-CGGYTACCTTGTTACGACTT-3’). Modified versions of these primers were designed with the following architectures:

#### Direct DBCO modification

A 5′-DBCO or 5′-azide modification was appended directly to the 1492R primer for initial SPAAC moiety screening.

#### Overhang configuration

The barcode flanking sequence (BFS) from the SQK-RBK114 MuA cargo (5′-GCTTGGGTGTTTAACC-3’) was appended upstream of the 1492R sequence, separated by a 12-carbon spacer (Sp12) to block polymerase extension. A 5′-DBCO modification was placed at the terminus of the BFS overhang, yielding: 5′-/DBCO/GCTTGGGTGTTTAACC/Sp12/CGGYTACCTTGTTACGACTT-3’.

#### Universal rapid oligonucleotide

A universal modified oligonucleotide was constructed by replacing the target-specific primer sequence with an M13 Forward universal sequence (5′-GTAAAACGACGGCCAGTG-3’), yielding: 5′-/DBCO/GCTTGGGTGTTTAACC/Sp12/GTAAAACGACGGCCAGTG-3’.

Corresponding target-specific primers (27F) carried the M13F sequence as a 5′ tail: 5′-GTAAAACGACGGCCAGTG-barcode-AGRGTTYGATYMTGGCTCAG-3’. This universal oligonucleotide was used in two configurations: (1) as a third primer in a single-pot reaction alongside unmodified 27F and 1492R, and (2) in a second-round PCR using purified unmodified first-round product as template.

#### RLB-compatible primers

For comparison with the ONT Rapid PCR Barcoding Kit (SQK-RPB114.24), target-specific primers were appended with RLB-compatible tail sequences: RLB-27F (5′-CGTTTTTCGTGCGCCGCTTCAGRGTTYGATYMTGGCTCAG-3’) and RLB-1492R (5′-CGTTTTTCGTGCGCCGCTTCCGGYTACCTTGTTACGACTT-3’).

All commercially modified oligonucleotides were synthesized by Integrated DNA Technologies (IDT) or Eurogentec.

### In-house DBCO conjugation via TFP ester

To further explore flexibility, 5′-amino modified universal oligonucleotides were ordered from IDT and conjugated to DBCO using the EZ-Link TFP Ester-PEG4-DBCO labeling kit (Thermo Fisher Scientific). Reactive primary amine oligonucleotides were dissolved in 0.5 M sodium carbonate buffer (pH 8.5) at 1 mM. TFP Ester-PEG4-DBCO was dissolved in anhydrous DMSO at 10 mM. Equal volumes of oligonucleotide and labeling reagent were mixed (10:1 molar excess of TFP ester to oligonucleotide), incubated for one hour at room temperature, and then overnight at 4 °C. All incubations were protected from light. Reactions were diluted to 150 μM oligonucleotide concentration with 10 mM Tris-EDTA (TE) buffer for easier handling and to quench unreacted ester.

Using this protocol, oligonucleotides with spacer lengths of C12, C9, C3, and C0 (no spacer) were synthesized and labeled to assess the impact of spacer length on RA binding efficiency.

### Post-labeling purification

Amersham MicroSpin G-25 purification columns (Cytiva, 27532501) were pre-equilibrated by centrifugation for 1 minute at 735 x g. Twenty-five microliters of labeling mixture was loaded onto the column (a reduced volume chosen to increase purity as per manufacturer’s instructions). Columns were placed in collection tubes and centrifuged for 2 minutes at 735 x g to elute purified oligonucleotides.

### Library preparation and sequencing

Following PCR and bead cleanup, libraries were prepared by incubating purified amplicons with ONT Rapid Adapter (RA) for 5 minutes at room temperature. No enzymatic steps (end-repair, ligation, or transposase treatment) were performed. Prepared libraries were loaded onto primed PromethION flow cells (R10.4.1 chemistry) according to the manufacturer’s instructions.

### Flow cell reuse protocol

All 11 experimental conditions were sequenced on the same PromethION flow cell (PBK58990) to enable direct comparison. This new flow cell was primed according to the manufacturer’s instructions prior to the first sequencing run. Following each sequencing iteration, the flow cell was washed using the Flow Cell Wash Kit (EXP-WSH-004, ONT). After washing, the flow cell was loaded with storage buffer and sequenced for 0.1 hours (~6 minutes) to confirm no carryover between conditions. Sequencing runs were 0.25 hours per condition.

### Data analysis

#### Pore occupation

Pore occupation was defined as the fraction of pores actively sequencing strand DNA, divided by the total pores in sequencing, adapter, or available states, excluding inactive, unclassifiable, or permanently blocked pores. The adapter fraction was calculated separately using the same denominator. Measurements were taken at steady state (t = 10-15 minutes) to ensure stable readings.

#### Read normalization

The total number of reads passing the default quality filter (Q > 10) and within the expected amplicon size range (1,400-1,700 bp) were normalized to the number of available pores measured at t = 5 minutes per run, or the initial within run pore scan. This window accommodates the ~100 bp increase in product length for configurations that add tails to the target-specific primers (e.g., M13 universal tail, RLB tails), shifting the amplicon size from ~1,500 bp to ~1,600 bp and species variation around the average amplicon size. This normalization accounts for the progressive reduction in total pore count due to successive flow cell handling and time-dependent pore loss.

## RESULTS

The primer configurations evaluated in this study are summarized in Figure 1, progressing from the simplest direct DBCO attachment to the most flexible and cost effective two-PCR universal workflow. All configurations employ an asymmetric design in which the DBCO modification is restricted to a single primer, leaving the opposite primer available for barcoding.

**Figure 1.**
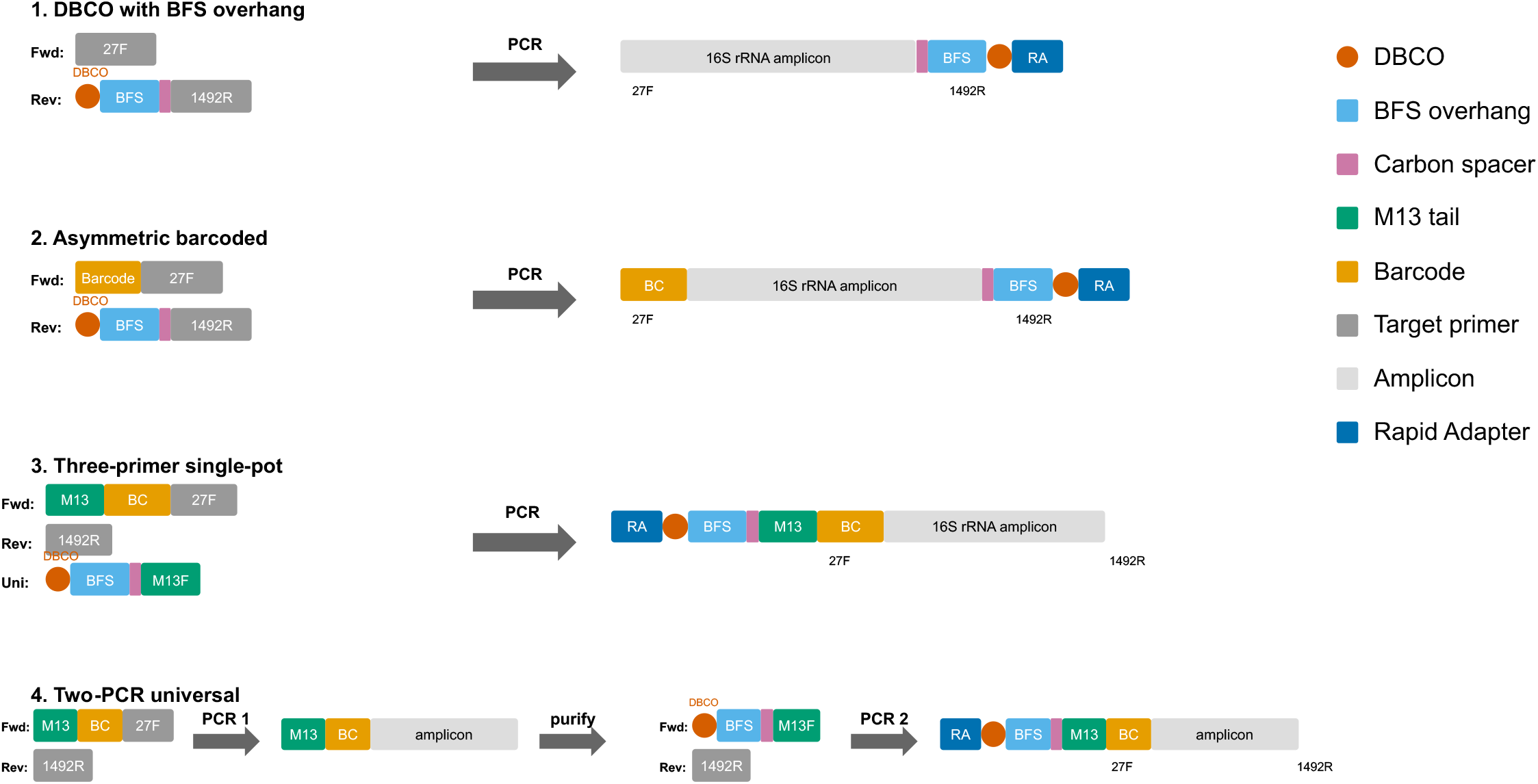
Primer architectures for DBCO-modified amplicon library preparation. Four configurations are shown, progressing from the simplest (direct DBCO attachment) to the most flexible (two-step PCR with universal DBCO oligonucleotide). Left: primer/oligonucleotide composition. Right: resulting amplicon structure after PCR, showing Rapid Adapter (RA) conjugation via SPAAC. All configurations are asymmetric, with DBCO restricted to one amplicon terminus. The carbon spacer blocks polymerase extension, generating the single-stranded BFS overhang required for efficient RA binding. Barcodes on the opposite primer enable sample multiplexing.

### SPAAC moiety selection: azide versus DBCO

SPAAC conjugation requires an azide and a strained cycloalkyne (or derivatives), but the polarity of the modification required on the amplicon was unknown. We therefore tested both available SPAAC moieties (5′-azide and 5′-DBCO) appended to the 1492R primer, paired with unmodified 27F. An unmodified 27F/1492R reaction served as a positive control for amplification. As shown in Supplementary Figure 1, neither modification affected PCR amplification, yielding post-cleanup concentrations of 27.2 ng/μL (azide), 16.3 ng/μL (DBCO), and 20.6 ng/μL (unmodified control).

After incubation with RA and loading onto the PromethION flow cell, the 5′-azide condition showed 7.3% sequencing pore occupation with 31.6% of pores in the adapter state, (Figure 2a) but produced zero reads in the 1,400-1,700 bp range, indicating that while RA entered pores, it did not carry any amplicon DNA (Figure 2b). While the 5′-DBCO condition achieved 11.7% sequencing occupation with an even higher adapter fraction (47.5%), similar to the azide modification. However, it successfully generated 1,103 reads of the expected length (Figure 2b). The high adapter-to-sequencing ratio in both conditions suggests that amplicons were not efficiently ligated to RA even if the adapter was present in sufficient amounts and successfully captured by the pores. Nonetheless, the DBCO condition produced reads while the azide did not, confirming that DBCO is a preferred SPAAC partner required for amplicons to bind the ONT Rapid Adapter and that the modification survives PCR thermocycling.

**Figure 2.**
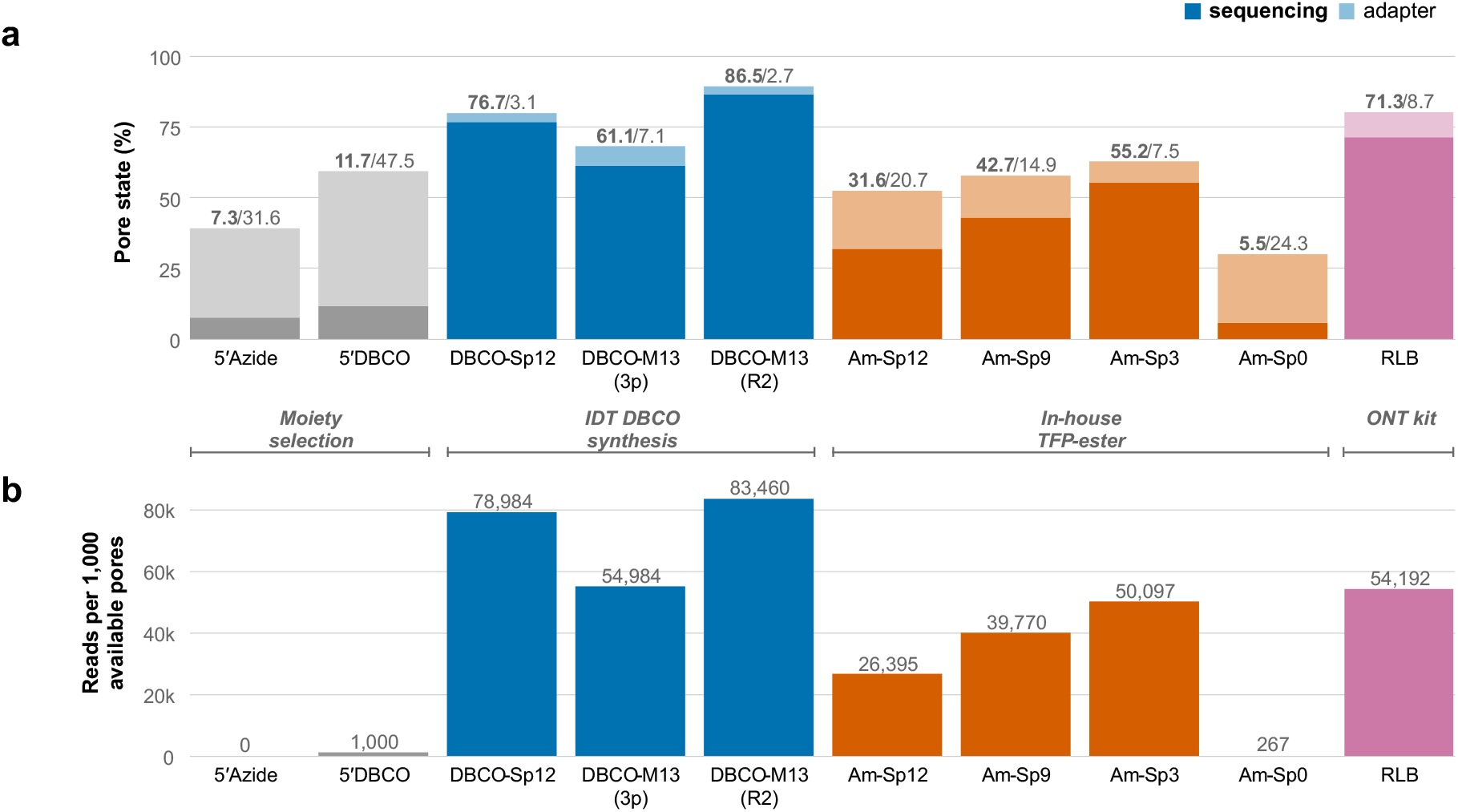
Pore occupation, adapter binding, and sequencing yield across 11 DBCO-primer configurations. All conditions were sequenced on the same PromethION flow cell (PBK58990) with washes between runs. (a) Stacked bars showing the fraction of pores actively sequencing (solid) and in adapter state (lighter shade), as a percentage of total sequencing + adapter + available pores at steady state (t = 10-15 min). Values above bars indicate sequencing%/adapter%. (b) Reads per 1,000 available pores (Q > 10, 1,400-1,700 bp), normalized to available pores at t = 5 min. The two-PCR workflow with commercially synthesized DBCO achieved the highest sequencing occupation (86.5%) with the lowest adapter fraction (2.7%), while the Amino-Sp0 condition (5.5% sequencing, 24.3% adapter) confirmed the requirement for a polymerase extension blocker.

### Enhanced adapter binding via a 5′ overhang (Configurations 1 & 2)

We hypothesized that the low read yield from direct 5′-DBCO attachment reflected inefficient initial association between the DBCO-modified amplicon terminus and the RA. We reasoned that a single-stranded 5′ overhang might be necessary to enhance binding, analogous to how complementary overhangs facilitate Native Adapter ligation in ONT ligation kits (Figure 3a) over blunt end ligation. We previously reported the use of MuA inserted cargo as a target to amplify phage genomes by PCR (Fox et al., 2025) and noticed that the sequence upstream of the barcode region was blocked from polymerase amplification. We appended the barcode flanking sequence (BFS; 5′-GCTTGGGTGTTTAACC-3’) (Oxford Nanopore, 2024) from the SQK-RBK114 MuA cargo upstream of a long 12-carbon spacer (Sp12) and the 1492R sequence, with the same partially successful 5′-DBCO at the overhang terminus. The carbon spacer serves as a polymerase extension blocker, ensuring the BFS remains single-stranded after PCR leading to an overhang. A 12-carbon spacer was initially selected at this length to ensure full blocking of the highly processive SuperFi II polymerase used in the reactions.

**Figure 3.**
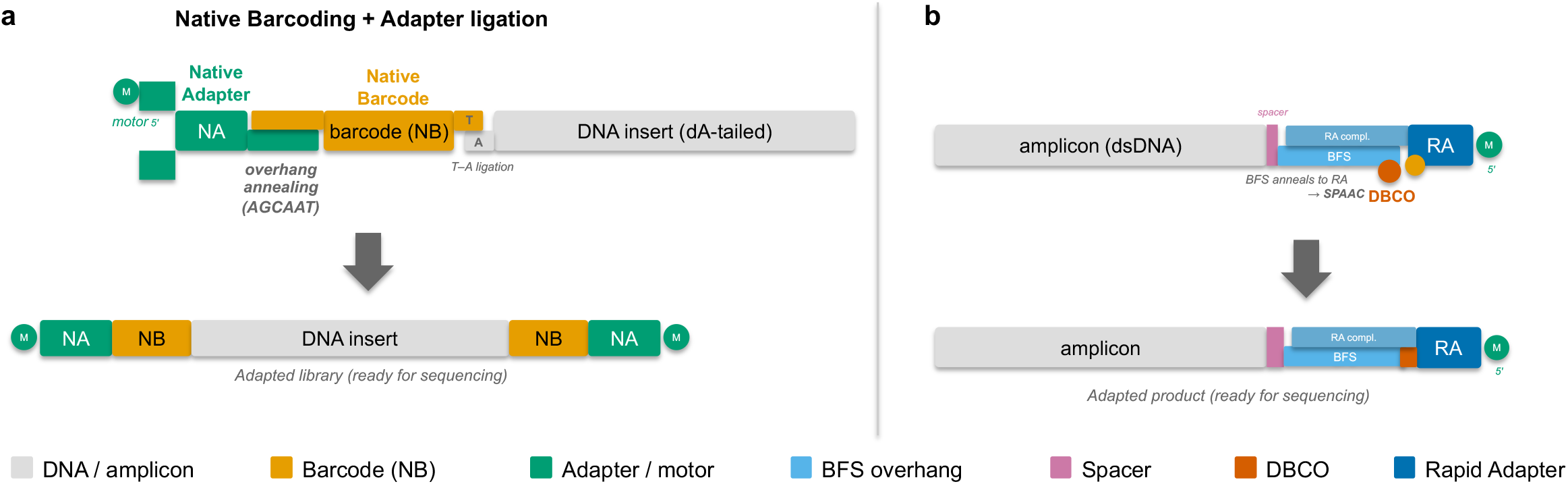
Proposed model of Rapid Adapter binding enhanced by a 5′ single-stranded overhang. (a) In the standard ONT workflow, Native Barcodes ligate to dA-tailed DNA, and the Native Adapter binds to the barcode via a complementary overhang. (b) In the DBCO-primer approach, the carbon spacer blocks polymerase extension, generating a single-stranded 5′ BFS overhang after PCR. This overhang anneals to a complementary sequence on the opposite strand of the Rapid Adapter, positioning the DBCO (on the amplicon) in close proximity for spontaneous SPAAC conjugation without enzymatic ligation.

In this model (Figure 3b), the single-stranded overhang produced by polymerase blocking enhances initial binding of the amplicon to the RA by annealing to complementary sequences on the opposite strand of the adapter, positioning the click chemistry groups in close proximity for efficient SPAAC conjugation.

To test whether the addition of spacer elements and the overhang would affect PCR and subsequent ligation efficiency, we compared reactions from unmodified products, the partially successful DBCO modified products and the DBCO modified product with an induced overhang. PCR yields were 19.9 ng/μL (unmodified), 29.2 ng/μL (5′-DBCO only), and 8.94 ng/μL (5′-DBCO-BFS-Sp12) (Supplementary Figure 2). Even with a lower final library concentration, the addition of the BFS overhang dramatically improved adapter binding efficiency as evidenced by productive translocation: sequencing pore occupation increased from 11.7% to 76.7%, while the adapter fraction dropped from 47.5% to 3.1%, indicating that the vast majority of pore-captured molecules were now full amplicons ligated adapters (Figure 2a; left blue bar). Sequencing yield increased from 1,000 to 78,984 reads per 1,000 available pores, a 79-fold improvement (Figure 2b; left blue bar). This result confirmed that a single-stranded overhang is critical for efficient RA conjugation, consistent with the proposed binding model (Figure 3b). This modification to target primers allows for direct sequencing of a sample post PCR-cleanup (Configuration 1). Additionally, since efficient binding was seen using a single end modified amplicon, this asymmetric modification also allows for multiplexing by barcoding the opposite oligonucleotide without the additional synthesis cost of DBCO and spacer modifications (Configuration 2).

### Universal rapid oligonucleotide configurations

To enable a single modified oligonucleotide to serve multiple amplicon targets, we designed a universal DBCO-modified primer bearing an M13 Forward tail instead of a target-specific sequence. This universal oligonucleotide (5′-/DBCO/GCTTGGGTGTTTAACC/Sp12/GTAAAACGACGGCCAGTG-3’) was tested in two configurations.

#### Three-primer single-pot reaction (Configuration 3)

The M13-tailed barcoded 27F primer (5′-GTAAAACGACGGCCAGTG-barcode-27F-3’), unmodified 1492R, and the universal DBCO-M13 oligonucleotide were combined in a single PCR. In early cycles, the M13-tailed 27F and 1492R amplify the target; in later cycles, the universal DBCO-M13 oligonucleotide anneals to the M13 tail and extends, incorporating the BFS overhang and DBCO modification. This reaction yielded 19.7 ng/μL (Supplementary Figure 3), compared to 14.4 ng/μL for the unmodified control and 10.2 ng/μL for the fully modified 1492R oligonucleotide. Sequencing pore occupation reached 61.1% with an adapter fraction of 7.1% and 54,984 reads per 1,000 available pores (Figure 2; middle blue bars). Although less efficient than the direct overhang configuration, this approach produced substantial sequencing data while requiring only a single modified oligonucleotide that can be reused across any amplicon target.

#### Two-PCR workflow (Configuration 4)

A first-round PCR with M13-tailed 27F and unmodified 1492R primers (14.4 ng/μL yield) was followed by a 15-cycle second-round PCR using 1 μL of purified first-round product as template, with the universal DBCO-M13 oligonucleotide and unmodified 1492R. The M13 tail from PCR1 provides the annealing site for the universal DBCO-M13 primer in PCR2. The second-round PCR yielded 18.5 ng/μL (Supplementary Figure 4). This configuration achieved the highest performance of all conditions tested: 86.5% sequencing pore occupation with the lowest adapter fraction (2.7%) and 83,460 reads per 1,000 available pores (Figure 2; right blue bars). The low adapter fraction indicates that nearly all pore-captured molecules were amplicon ligated RA, consistent with efficient DBCO+overhang mediated conjugation.

**Figure 4.**
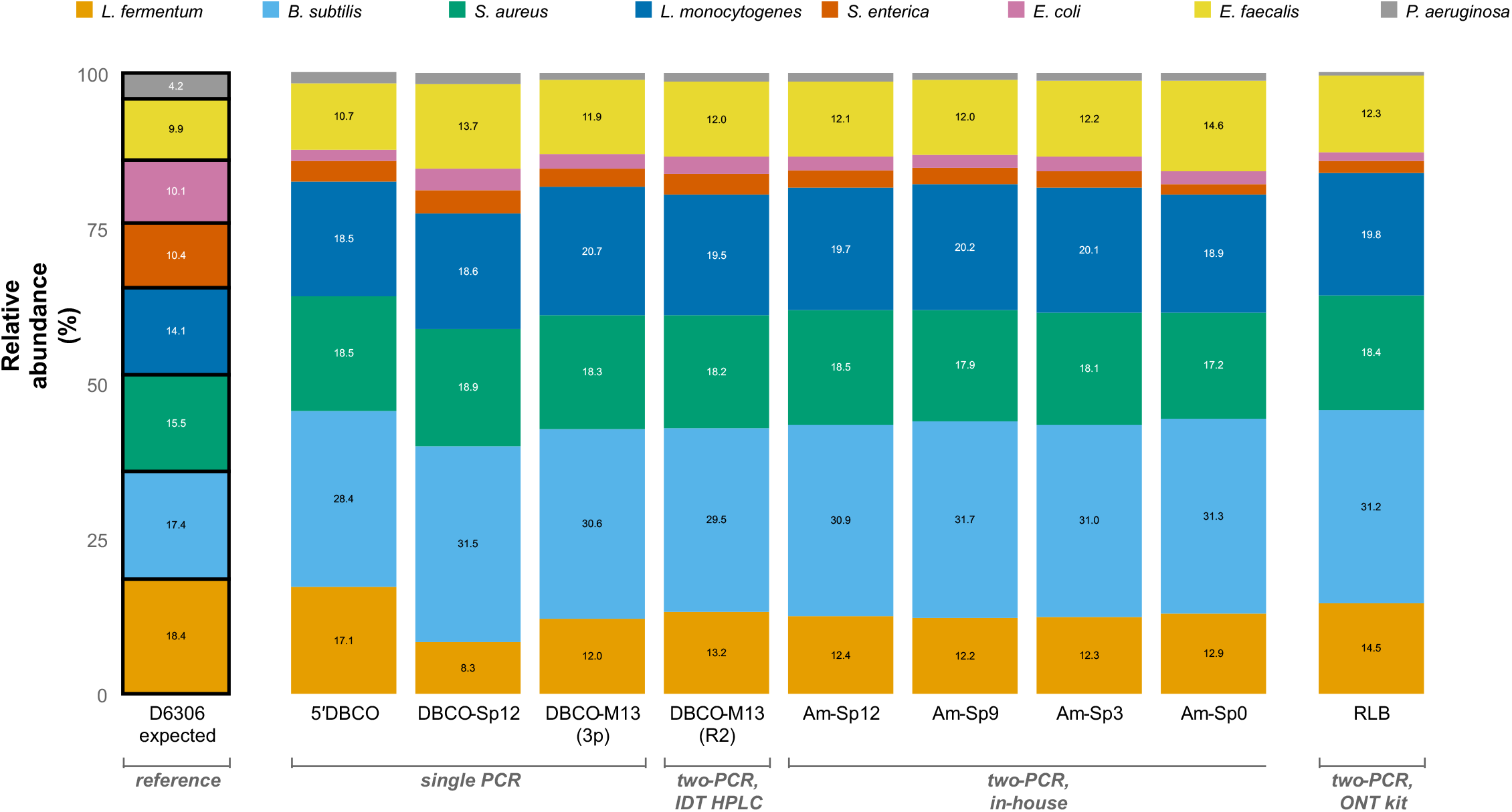
16S rRNA gene community composition across experimental conditions compared to the expected ZymoBIOMICS D6306 standard. Stacked bars show the relative abundance (%) of each of the eight bacterial species in the mock community, with percentages labeled inside each segment. The expected composition (leftmost bar, outlined in black) is derived from the D6306 theoretical abundances. Conditions are grouped by PCR workflow and purification method (brackets below). The universal M13-based configurations (two-PCR) produced highly consistent profiles, while the direct DBCO-Sp12 configuration (single PCR) showed a modest shift in *L. fermentum* (8% vs. 12-13%). The systematic over-representation of Gram-positive species and under-representation of Gram-negative species is shared across all conditions including the RLB ONT kit control, confirming this is a property of the 27F/1492R primer pair, not the adapter chemistry.

### In-house DBCO conjugation and spacer length optimization

To reduce per-oligonucleotide costs, we synthesized 5′-amino modified universal oligonucleotides and conjugated DBCO via TFP ester chemistry. Simultaneously, we investigated the effect of spacer length on RA binding by comparing 12-carbon, 9-carbon, 3-carbon, and 0-carbon (no spacer) configurations, testing whether the spacer or the BFS overhang was primarily responsible for enhanced binding.

PCR yields using the in-house conjugated oligonucleotides were: IDT-DBCO-Sp12 (commercial control) 13.8 ng/μL, Amino-Sp12 4.96 ng/μL, Amino-Sp9 4.14 ng/μL, Amino-Sp3 9.24 ng/μL, and Amino-Sp0 14.1 ng/μL (Supplementary Figure 5). All in-house conjugated oligonucleotides with spacers (Sp12, Sp9, Sp3) were purified using G-25 spin columns.

Sequencing results revealed a clear spacer-length threshold (Figure 2; orange bars). Sp12, Sp9, and Sp3 all produced functional libraries with sequencing pore occupations of 31.6%, 42.7%, and 55.2%, adapter fractions of 20.7%, 14.9%, and 7.5%, and read yields of 26,395, 39,770, and 50,097 reads per 1,000 available pores, respectively with variations in efficiency likely stemming from differences in purification and handling. The C0 (no spacer) configuration however, in which DBCO was attached directly to the terminal nucleotide without a polymerase blocker, showed only 5.5% sequencing occupation with 24.3% adapter fraction and 267 reads per 1,000 available pores, a near-complete loss of productive translocation comparable to the initial direct DBCO attachment without an overhang. These results demonstrate that a polymerase extension blocker of at least C3 is essential for generating the single-stranded overhang required for efficient RA binding, while spacer lengths beyond C3 are unlikely to confer additional benefit.

The lower overall performance of in-house conjugated oligonucleotides relative to the commercially synthesized IDT-DBCO version (31.6-55.2% vs. 86.5% pore occupation) likely reflects incomplete labeling and the inability of G-25 columns to separate labeled from unlabeled oligonucleotides, resulting in a mixture of DBCO-bearing and unreactive amplicons that leads to a lower effective concentration of sequencable amplicons.

### Compatibility with ONT Rapid PCR Barcoding primers

To benchmark DBCO-primer performance against a standard ONT workflow, we adapted the universal primer protocol to the ONT Rapid PCR Barcoding Kit (SQK-RPB114.24) four-primer approach. First-round PCR with RLB-compatible tailed primers yielded 44.8 ng/μL and was diluted to 14.4 ng/μL to be comparable to our M13F-tailed amplicon pool used above. 1 μL was carried forward into a second-round PCR with 1 μL of RLB barcoded primer and yielding 13.7 ng/μL post PCR-cleanup.

The RLB condition achieved 71.3% sequencing pore occupation with 8.7% adapter fraction and 54,192 reads per 1,000 available pores (Figure 2; pink bar). This places the RLB kit between the three-primer single-pot DBCO configuration (61.1% pore occupation) and the two-PCR DBCO configuration (86.5% pore occupation), demonstrating that the DBCO-primer approach can match or exceed standard kit performance while offering substantially greater flexibility and lower per-reaction costs.

### Community composition analysis

To verify that the DBCO-primer library preparation does not introduce bias, especially when using a single vs. double PCR configuration, we compared the 16S rRNA gene community composition from each configuration against the expected composition of the ZymoBIOMICS Microbial Community Standard (D6306). Reads passing quality (Q > 10) and length (1,400-1,700 bp) filters were aligned to a reference database containing the 16S rRNA gene sequences of the eight bacterial species in the standard using minimap2, and relative abundances were calculated from primary alignments.

All configurations that produced sufficient reads (n = 9, excluding the 5’-azide negative control) yielded broadly concordant community profiles sharing the same systematic bias pattern (Figure 4). The mean L1 distance (sum of absolute differences in relative abundance across all eight species) from the expected composition was 47.2 across all conditions. The same directional bias was observed in every condition: over-representation of Gram-positive species, particularly *Bacillus subtilis* (observed 28-32% vs. expected 17.4%) and *Listeria monocytogenes* (18-21% vs. 14.1%), and under-representation of Gram-negative species, particularly *Salmonella enterica* (2-4% vs. 10.4%), *Escherichia coli* (1.8-3.4% vs. 10.1%), and *Pseudomonas aeruginosa* (0.6-1.8% vs. 4.2%). The RLB four-primer ONT kit condition showed an identical bias pattern (L1 = 49.4), confirming that this is a property of the 27F/1492R primer pair rather than the adapter attachment chemistry.

Among the universal M13-based configurations (Configurations 3-4 and all in-house spacer variants), per-species abundances were highly consistent, generally varying by less than 2% across conditions regardless of spacer length, synthesis method, or number of PCR rounds. The direct DBCO-Sp12-1492R configuration (Configuration 2) showed a modest compositional shift, with lower *Lactobacillus fermentum* (8.3% vs. 12-13% in M13-based configurations) and higher *Enterococcus faecalis* (13.7% vs. ~12%), which may reflect the different primer architecture (target-specific vs. universal). These differences do not alter the overall conclusion that the DBCO click chemistry adapter attachment itself has no impact on taxonomic bias.

## DISCUSSION

We have demonstrated that DBCO-modified PCR primers enable direct, transposase-free conjugation of nanopore sequencing adapters via SPAAC click chemistry, reducing amplicon library preparation to a single post-PCR pipetting step. The approach achieved up to 86.5% pore occupation on PromethION, exceeding standard rapid kit performance (71.3% for the RLB four-primer protocol), with lower free adapter levels, simpler laboratory workflow, and a significant reduction in per-sample reagent costs as described below. At a 250nmol synthesis scale, IDT offers a guaranteed 6.5nmol minimal yield for the 34bp universal oligo, sufficient for 650 PCR reactions at 0.2 μM in 50 μL but likely to be sufficient for more reactions as IDT synthesis often results in more than the minimum yield. In contrast, combining all barcodes across the Rapid PCR Barcoding Kit yields only 0.09nmols. Including HPLC purification, DBCO and spacer modifications, the IDT list price at 623$ CAD while both the Rapid PCR Barcoding kit and Microbial Amplicon Barcoding Kit are listed at 1,180$ CAD each. Additional auxiliary buffers and RA are available for 170$ CAD and 490$ CAD respectively.

The requirement for a carbon spacer between the BFS and the primer target sequence prevents polymerase extension and generates a 5′ overhang that likely contributes to more efficient initial binding to RA by maintaining the reactive moieties in close proximity. Our data demonstrate that spacers as short as C3 are sufficient, while direct attachment of DBCO to the terminal nucleotide without a spacer drastically reduces conjugation efficiency (5.5% pore occupation, 267 reads per 1,000 pores). That C3, C9, and C12 spacers perform similarly suggests that beyond acting as a polymerase extension blocker, additional spacer length confers no advantage.

Community composition analysis confirmed that the DBCO-primer approach does not introduce obvious sequence-dependent taxonomic bias. All configurations produced 16S profiles sharing the same systematic bias pattern as the RLB kit control (Figure 4). The universal M13-based configurations were highly consistent (per-species variation < 2%), while the direct DBCO-Sp12-1492R configuration showed a modest compositional shift (e.g., *L. fermentum* 8.3% vs. 12-13%), likely reflecting the different primer architecture rather than the adapter chemistry. The observed Gram-positive over-representation is a well-documented property of the 27F/1492R primer pair (Matsuo et al., 2021; Nygaard et al., 2020) and is unrelated to the adapter attachment chemistry.

Two complementary synthesis routes provide flexibility for different laboratory settings. Commercial synthesis of DBCO-modified oligonucleotides offers convenience but is currently available from a limited number of suppliers and incurs a modification surcharge. The in-house route, purchasing standard 5′-amino oligonucleotides at a lower surcharge and conjugating DBCO-PEG4-TFP ester can be more cost-effective at scale because a single vial of TFP ester reagent can modify dozens of oligonucleotides, but the economics of purchasing additional reagents versus a larger number of modified oligonucleotides remain to be determined on a per lab basis. TFP esters offer superior aqueous stability compared to NHS esters, which hydrolyze rapidly at physiological pH, improving conjugation yield and reproducibility (Thermo Fisher, 2023). The commercial synthesis, however, offers greater purity through HPLC purification options, whereas in-house synthesis purification options are limited to centrifugation column kits unless core HPLC facilities are available. In addition to potentially leaving more unreacted DBCO labeling molecules behind, centrifugation columns also do not segregate labeled from unlabeled oligonucleotides, leading to a potentially reduced effective RA binding capacity for equimolar final amplicon pools. PAGE purification remains a potentially useful choice where available as it has the ability to discriminate labeled from unlabeled oligonucleotides.

The asymmetric single-modification configuration merits particular attention for cost-sensitive applications with a balanced time cost. By restricting the DBCO modification to one primer per pair and using inexpensive unmodified barcoded primers on the reverse strand, this design requires a single modified oligonucleotide per project and can be paired with a nearly unlimited number of barcodes on the opposite primer. A 5′ 20 bp padding on barcoded primers is however essential to ensure barcode sequences remain within the basecaller’s high-confidence region due to 3′-terminal signal degradation during ONT sequencing, slightly increasing synthesis cost. Demultiplexing with Dorado’s custom barcode arrangement system only requires a few bases upstream with a full sequence downstream (PCR target), though we note that not all IUPAC ambiguity codes are supported in Dorado flanking sequence definitions, which can cause issues for users designing panels with highly degenerate primers. We recommend padding an additional known sequence between the barcode and the target region as a potential mitigating strategy. This limitation can likely be circumvented by future updates to the Dorado source code or compilation of a custom version capable of handling further degenerate codes, as partial implementation for “N” and “M” bases already exists.

The Microbial Amplicon Barcoding Kit (SQK-MAB114.24), released in September 2025 as an Early Access product, addresses primer bias by offering redesigned 16S sequences with improved inclusivity and adding ITS primers for fungal profiling (Oxford Nanopore, 2025). Crucially, however, it introduces a multi-step post-PCR protocol that substantially increases hands-on time: amplicons must undergo barcode ligation, proteinase K digestion, heat inactivation, bead cleanup, and finally RA attachment. This series of incubation and enzymatic processing steps extends total library preparation time well beyond the older 16S kit’s simpler workflow, partially negating the convenience that bundled kits are designed to provide, and the kit retains the 24-barcode ceiling. By contrast, the DBCO-primer approach can reduce the entire post-PCR protocol to a single adapter addition at room temperature, depending on the implementation.

The Rapid PCR Barcoding Kit (SQK-RPB114.24) provides an alternative four-primer barcoding approach with 24 barcodes, using ONT-tailed primers for a two-step PCR workflow. While conceptually similar to our universal DBCO-primer configurations (Configurations 3 and 4), the kit provides only a small volume of 10 μM barcoded primers per barcode position, a quantity that is rapidly exhausted in routine use, particularly for studies requiring optimization replicates or high-throughput screening. In contrast, DBCO-modified primers synthesized commercially or via DBCO-TFP ester conjugation to 5′-amino oligonucleotides frequently yield 50-200 μL at 100 μM (well above guaranteed minimum yield), sufficient for thousands of PCR reactions per synthesis.

Relative to the general-purpose library preparation kits, the advantages are equally clear. Ligation sequencing (SQK-LSK114) requires end-repair, dA-tailing, adapter ligation, and two bead cleanups (~75 minutes of hands-on time). Rapid barcoding (SQK-RBK114) requires transposase incubation (2 min, 30 °C), heat inactivation (2 min, 80 °C), pooling, bead cleanup, and adapter addition, while truncating 10-20 bp from amplicon ends but most importantly, fragmenting the input amplicon and leading to unlinked 5′ and 3′ sequences. The DBCO-primer workflow eliminates all enzymatic steps: after PCR and a standard bead cleanup, the RA is added directly to the purified amplicon, and the library is loaded onto the flow cell following a short incubation at room temperature.

The higher pore occupation achieved by DBCO-primer libraries (up to 86.5% vs. 71.3% for the standard RLB kit; lower for in-house DBCO-TFP conjugation at 31.6-55.2%) likely reflects a combination of the shorter timeline between synthesis and use compared to commercial kits involving warehousing and delivery, and a more selective purification post-labeling. Indeed, while the Cytiva G-25 resin can remove excess labeling reagents and salts, the purification does not discriminate between labeled and unlabeled oligonucleotides, unlike the HPLC purification step in commercial synthesis reactions. The lower free adapter signal further supports this interpretation: fewer unreacted adapter molecules in the library means fewer adapter-only events that occupy pores without generating useful data.

The universal primer configurations (Configurations 3 and 4) are particularly relevant for large amplicon panels, such as those used in antimicrobial resistance profiling or viral surveillance. Rather than synthesizing DBCO-modified versions of every target-specific primer pair, a single universal DBCO-primer can serve all targets in the panel. The two-PCR workflow provides the cheapest implementation, while the three-primer single-pot variant offers closed-tube convenience at the cost of requiring careful primer concentration optimization to ensure efficient handoff between target-specific and universal amplification phases. Importantly, these universal configurations are not subject to the primer volume constraints of the Rapid PCR Barcoding Kit, because the universal DBCO-primer is a single oligonucleotide synthesis that can be used for thousands of reactions. In the case of the 2-step PCR, expensive modified oligonucleotides are only used **once** per sequencing run resulting in sufficient reagents (at minimum guaranteed yield) for 650 × 96+ [# barcodes per pool] = 62,400+ samples.

A particularly compelling application of the universal DBCO-primer approach is in UMI-based amplicon sequencing, where it could substantially simplify the post-PCR library preparation bottleneck. UMI workflows have transformed nanopore amplicon accuracy: the ssUMI method developed by Karst et al. (Karst et al., 2021) achieves consensus error rates below 0.005% (>Q43) by tagging individual template molecules with dual 18 bp UMIs during a low-cycle first PCR, then re-amplifying with universal primers in a second PCR before generating consensus sequences from UMI-binned subreads. (Lin et al., 2024) adapted this workflow for high-throughput full-length 16S rRNA profiling on R10.4.1 chemistry, achieving 99.99% accuracy at just 3x subread coverage per UMI bin, surpassing Illumina, and generating zero false-positive ASVs from mock communities.

All published UMI-nanopore workflows share the same post-PCR bottleneck: after the second-round universal primer amplification, the amplicon library must undergo ONT’s ligation sequencing protocol (SQK-LSK114). This entails end-repair and dA-tailing with NEBNext Ultra II enzymes (5 min at 20 °C, 5 min at 65 °C), a bead cleanup, adapter ligation with Salt-T4 DNA Ligase in ONT Ligation Buffer (10 min at room temperature), and a second bead cleanup using Short Fragment Buffer, totaling approximately 75 minutes of hands-on time with three to four bead-based purification steps. Ligation is required rather than rapid (transposase) preparation because transposase fragmentation would destroy full-length amplicons and unlink their terminal UMIs. Each bead cleanup step recovers 80-90% of input, so cumulative losses across three cleanups could potentially reach 25-35% of the library. The enzymatic reagents (NEBNext end-repair module, Salt-T4 ligase, SQK-LSK114 kit) add approximately $110-115 per library preparation.

The DBCO-primer approach offers a direct solution to this bottleneck. In the ssUMI workflow, the second-round PCR already uses a universal verification primer (UVP) that anneals to synthetic adapter tails incorporated during UMI tagging. If this universal primer carried a 5′-DBCO modification with an appropriate carbon spacer, every amplicon emerging from PCR2 would bear DBCO groups on both 5′ termini. After a single bead cleanup to remove primers and dNTPs, RA could be added directly, eliminating the end-repair, dA-tailing, enzymatic ligation, and their associated intermediate cleanups entirely. The proposed workflow would compress from four enzymatic/purification operations spanning ~75 minutes to a single bead cleanup followed by a room-temperature click reaction (~25-30 minutes total, with only one cleanup step). The cost of the oligonucleotide modification would be offset by savings from eliminated NEB enzymes plus reduced bead consumption.

This application is particularly well-suited to the universal primer configurations described here because the UMI workflow already mandates a two-step PCR with a common second-round primer (Configuration 4). No modification to the UMI-tagging first PCR is required; only the universal primer in the second PCR needs the 5′-DBCO addition. The UMI sequences, target amplicon, and primer binding sites remain unchanged, meaning existing ssUMI bioinformatic pipelines (longread_umi, ssUMI) would function without modification. This represents a drop-in improvement to an established workflow rather than a re-engineering of the entire protocol.

Several limitations should be noted. First, this work does not include a comprehensive analysis of barcode crosstalk or demultiplexing error rates, as this was not the focus of the study and has been covered extensively in the literature (Srivathsan et al., 2024); systematic benchmarking against standard ONT barcoding kits under controlled crosstalk conditions remains to be performed. With the flexibility of these protocols, however, reusing the 24 bp ONT barcodes may be the simplest implementation. Second, a comprehensive study of the stability of the DBCO modification over time has not been performed. It is entirely possible that the lifespan of the modification will be less than the speed at which the oligonucleotide is consumed, leading to extra costs from wastage. In our experience, however, the modification appears stable for at least 2 years when stored at −20 °C, as we did not notice any drastic reduction in binding efficiency as reflected by low free adapter levels and high pore occupation over this period.

In conclusion, DBCO-modified PCR primers provide a general, low-cost, enzyme-free method for generating nanopore sequencing libraries from any PCR amplicon. The approach resolves almost every major limitation of the current ONT bundled amplicon kits: it eliminates primer sequence constraints and the associated taxonomic biases, removes the 24-barcode multiplexing ceiling, replaces multi-step enzymatic protocols with a single adapter addition, and provides primer quantities sufficient for thousands of reactions from a single synthesis. We anticipate that this method will be particularly valuable for field-deployable sequencing, resource-limited settings, and high-throughput amplicon surveillance programs where simplicity, cost, and speed are most important.

## DATA AVAILABILITY

Sequencing data have been deposited at NCBI under the BioProject PRJNA1454126.

## FUNDING DECLARATION

This research was supported by Project Grants from the Canadian Institutes of Health Research (CIHR) to PL and BJS. This research was additionally supported by a Postdoctoral Fellowship from the Canadian Institutes of Health Research to PL.

## Notes

### Competing Interest Statement

The authors have declared no competing interest.

