## Supplementary Information for "Dibenzocyclooctyne-modified PCR primers enable direct enzyme-free click chemistry ligation for custom nanopore amplicon sequencing"

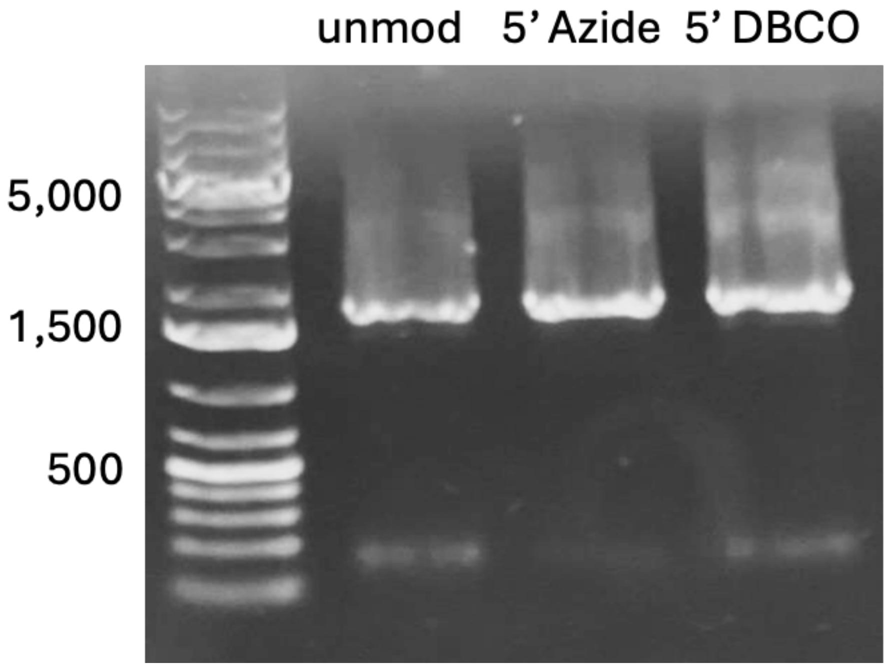


**Supplementary Figure 1. SPAAC moiety selection, gel electrophoresis of PCR products.** Agarose gel showing amplification of the full-length 16S rRNA gene (~1,500 bp) from ZymoBIOMICS mock community DNA. Lanes: DNA ladder; unmodified 27F/1492R; unmodified 27F/5′-Azide-1492R; unmodified 27F/5′-DBCO-1492R. All three conditions produced bands of the expected size, confirming that neither the azide nor DBCO 5′ modification interferes with PCR amplification.

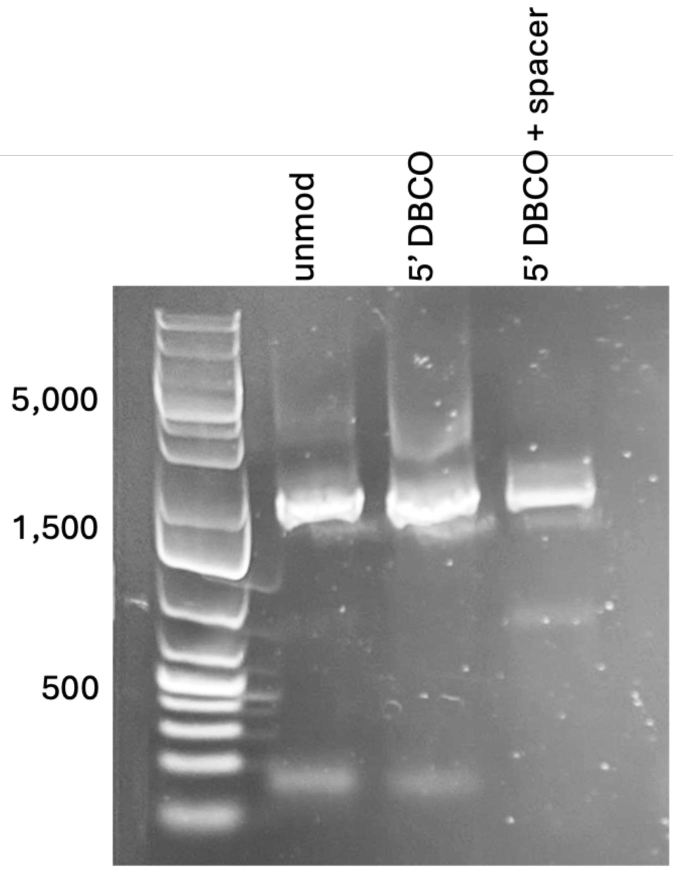


**Supplementary Figure 2. Overhang generation, gel electrophoresis of PCR products.** Agarose gel comparing amplification products with different 1492R primer designs. Lanes: DNA ladder; unmodified 27F/1492R; unmodified 27F/5′-DBCO-1492R; unmodified 27F/5′-DBCO-BFS-Sp12-1492R.

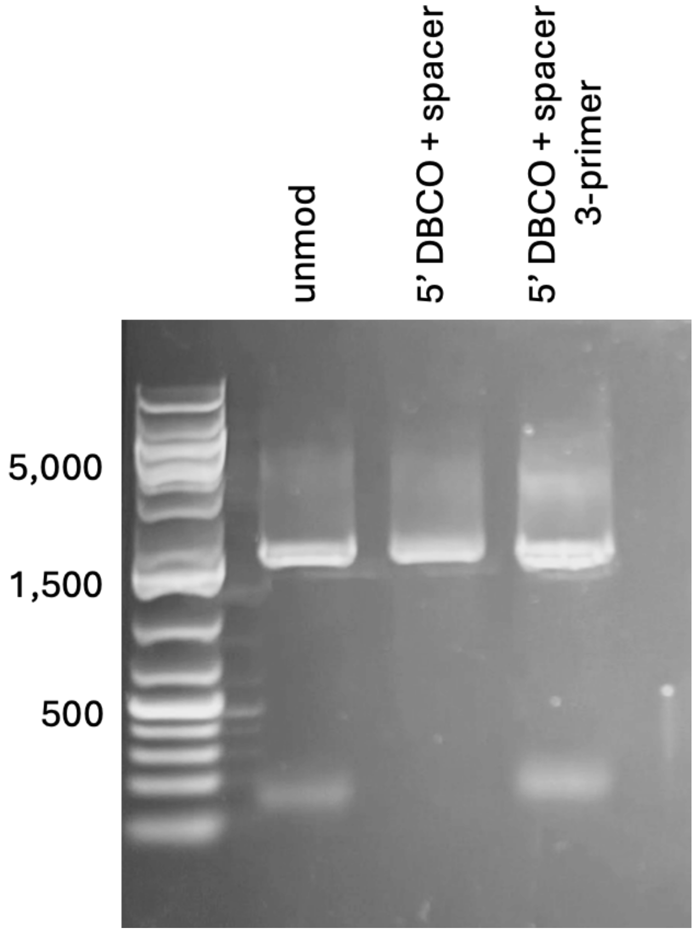


**Supplementary Figure 3. Universal rapid oligonucleotide single pot reaction.** Agarose gel showing amplification using the universal DBCO-M13 oligonucleotide. Lanes: DNA ladder; unmodified 27F/1492R; unmodified 27F/5′-DBCO-Sp12-1492R fully modified; three-primer reaction with DBCO-M13F + unmodified 27F + unmodified 1492R. All conditions produced bands of the expected size.

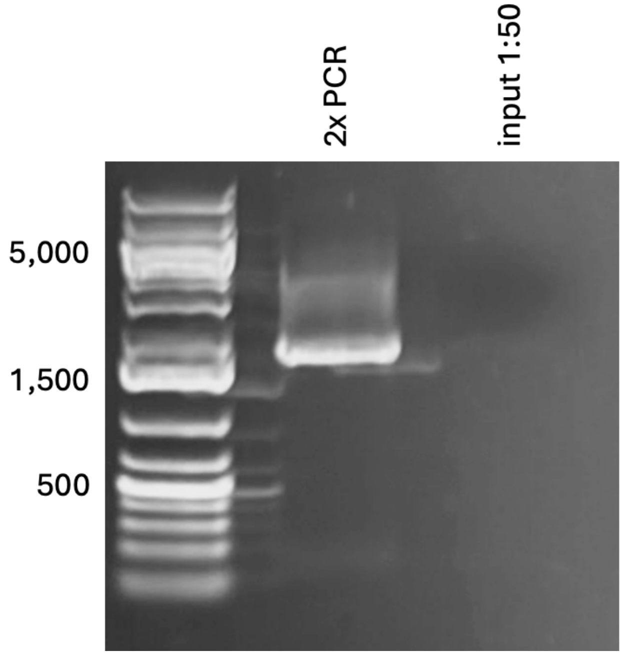


**Supplementary Figure 4. Second-round PCR with universal modified oligo.** Agarose gel showing the second-round PCR product amplified from 1 μL of purified unmodified first-round product (14.4 ng/μL stock) using the universal DBCO-M13 oligonucleotide and unmodified 1492R (15 cycles). Lane: DNA ladder; second-round PCR product; input purified first round product diluted to 1:50 to reflect PCR input concentrations.

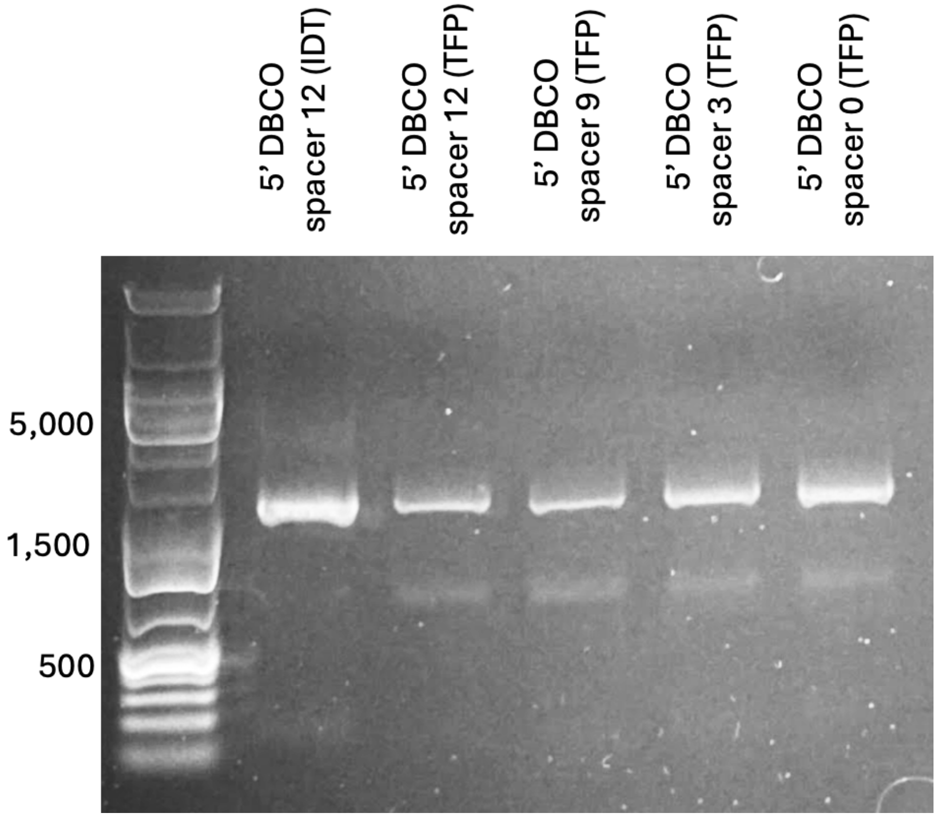


**Supplementary Figure 5. Spacer length comparison.** Agarose gel comparing amplification products using universal DBCO-M13 oligonucleotides with different spacer lengths, synthesized in-house via TFP ester conjugation to 5′-amino oligonucleotides. Lanes: DNA ladder; IDT-DBCO-Sp12 commercial control; 5′-Amino-Sp12; 5′-Amino-Sp9; 5′-Amino-Sp3; 5′-Amino-Sp0. All conditions show successful amplification.

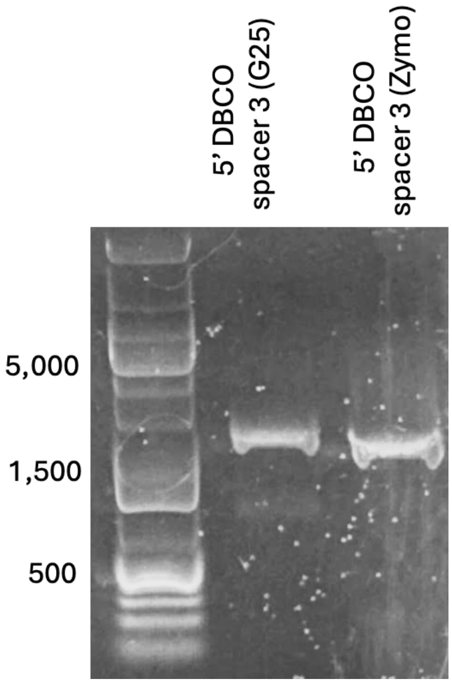


**Supplementary Figure 6. Post-labeling purification comparison.** Agarose gel comparing PCR products amplified using the 5′-Amino-Sp3 oligonucleotide purified by two different methods after DBCO-TFP ester conjugation. Lanes: DNA ladder; Amino-Sp3 purified with Cytiva G-25 resin; Amino-Sp3 purified with Zymo Oligo Clean and Concentrator.
